# Toward a Cell Culture Model of Portal Axis Lipid Handling

**DOI:** 10.1101/2022.11.30.518512

**Authors:** Joshua R. Cook, Peter H.R. Green, Rebecca A. Haeusler

## Abstract

The health risks posed by excessive visceral white adipose tissue (WAT) may arise from exposure to deleterious intestinal factors in transit through the portal system. These may include an altered bile acid (BA) pool – particularly with respect to 12α-hydroxylated BA (12-HBA), which we have previously demonstrated are closely associated with insulin resistance. However, given the complexity of the enterohepatic milieu, we have worked to develop a co-culture system that allows for controlled study of the interactions between human induced pluripotent stem cell (hiPSC) derived adipocytes and primary human intestinal organoids (hIO). We present preliminary data on the characterization of both cellular compartments and their interactions, particularly in the context of BA treatment. We have recapitulated the basic functionality of each tissue type and have found reciprocal, potentially important changes in gene expression in each. However, we also have encountered numerous technical challenges in the development of this co-culture system and provide methodological observations to assist others seeking to work with a co-culture system of this type.

## Introduction

Metabolic dysfunction in white adipose tissue (WAT) is thought to be an early event in the natural history of insulin resistance (IR) [1], and mesenteric (visceral) WAT seems to be particularly susceptible [2]. Although the outsize impact of visceral WAT on IR is multifactorial and not fully delineated, portal flux from the intestine appears to be an important factor [3, 4]. In addition to absorbed nutrients, potentially harmful substances can enter the splanchnic circulation from the intestinal epithelium, including gut flora or their byproducts, inflammatory mediators, or environmental toxins, among others [5]. Bile acids (BA) may represent another metabolic signal by which the intestine contributes to IR in visceral WAT [6-8]. In addition to their canonical facilitation of lipid absorption, BA influence systemic metabolism [7, 9] at least in part through transcriptional regulation of intestinal hormones such as fibroblast growth factor-19 (FGF-19) [10], peptide YY (PYY) [11], and glucagon-like peptide-1 (GLP-1) [12]. Similarly, we suspect that BA affect insulin action in visceral WAT by modulating intestinal secretion of these or other signals.

The signaling roles of BA have major implications for disorders of fat handling, including insulin-resistant states such as non-alcoholic fatty liver disease (NAFLD) and type 2 diabetes [9]. We have found a close and specific correlation in particular between insulin resistance (IR) and levels of 12α-hydroxylated bile acids (12-HBA), including cholic acid and its derivatives [6, 13]. The 12α hydroxylation reaction is catalyzed by CYP8B1, whose expression is downregulated by insulin through its inhibition of FoxO transcription factors [13]. It therefore follows that unrestrained FoxO activity in IR drives up CYP8B1 expression and 12-HBA synthesis [13]. However, 12-HBA may also play a causal role in the development of IR [14], thereby potentially establishing a pathologic cycle. As 12-HBA and non-12-HBA differentially affect gene expression in enterocytes [15], a shift in their balance may contribute to dysregulated metabolic coordination between intestine and WAT, in turn setting the stage for the development of IR.

It is difficult to parse out the contributions of individual cell types in as complicated a milieu as the small intestine; we therefore turn to cell-culture models as a starting point. In order to model the interactions of human small intestinal epithelial cells and adipocytes as faithfully as possible, we have taken a stem cell-based approach. We have designed a co-culture system in which human induced pluripotent stem cell (hiPSC)-derived adipocytes (iAC) are exposed to the basolateral surface of a polarized monolayer of human primary intestinal organoids (hIO) and then are treated with BA mixes of variable 12-HBA content. Here, we present preliminary characterizations of each cell type and their interactions as regards gene expression, insulin signaling, and lipid metabolism, as well as unbiased transcriptomics. Our primary objective is to report methodologic observations that may be useful for others seeking to work with such a research model.

## Materials & Methods

### Human primary intestinal organoid culture

Human primary intestinal organoids (hIO) were derived from biopsy specimens provided from endoscopies performed according to standard of care for medical indications, with informed consent of the donor, and with the approval of the Columbia University Human Research Protection Office. Biopsy specimens were collected in RPMI and were then washed twice with ice-cold PBS. Tissue was then minced into small pieces, rinsed again with ice-cold PBS, and then incubated for 30 min in Gentle Cell Dissociation Reagent (GCDR, StemCell Technologies). Tissue was then centrifuged and crypts were isolated by vigorous pipetting in ice-cold DMEM + 1% BSA followed by filtration through a sterile 70-μm cell strainer. Isolated crypts were then plated on 6-well tissue culture dishes in domes of growth factor-reduced, phenol red-free Matrigel (Corning) at 1000 crypts per dome. Domed crypts were grown in IntestiCult Organoid Growth Medium (StemCell Technologies) supplemented with Y-27632 (ROCK inhibitor) (10 μM, StemCell Technologies), penicillin-streptomycin 1% (Gibco), and Normocin (InvivoGen). Medium was changed every 2-3 days, excluding Y-27632. Organoids were passaged every 7-14 days by dissociating Matrigel domes in ice-cold GCDR, breaking up organoids by vigorous pipetting, and re-embedding organoids split 1:3-1:4 in new Matrigel domes in IntestiCult Organoid Growth Medium.

### Culture of hIO in 2D (monolayer)

Organoids in 3D culture were harvested by breaking up Matrigel domes in ice-cold GCDR. Pelleted organoids were then vigorously pipetted and incubated for 10 min in trypsin-EDTA (Gibco) to dissociate into single cells. Single cell pellets were washed in DMEM/F_12_, resuspended in culture medium (see below) containing penicillin-streptomycin, Normocin, and Y-27632 (10 μM), and plated on Corning Transwell inserts (polyester, 12 mm diameter, 0.22 μm pores) coated with a thin film of growth factor-reduced, phenol red-free Matrigel. Culture medium was applied as 0.5 mL in the upper chamber and 1.0 mL in the lower chamber and changed every other day; Y-27632 was omitted after the first medium change. Monolayers reached confluence within 4-7 days.

Two different culture media were tested for hIO growth and differentiation: one commercial (“SCT-ODM”: IntestiCult Organoid Differentiation Medium, StemCell Technologies) and one homemade cocktail (“CCM”: Crypt Culture Medium) formulated according to a recipe developed by another group [16]. The commercial medium was supplemented with GlutaMAX (Gibco). The homemade cocktail was prepared in a base medium of advanced DMEM/F_12_ containing HEPES 15 mM, GlutaMAX, penicillin-streptomycin, and Normocin, and contains N2 supplement (ThermoFisher, 1X), B27 supplement (ThermoFisher, 1X), *N*-acetylcysteine 1 mM (Sigma-Aldrich), epidermal growth factor (50 ng mL^-1^, ThermoFisher), Noggin (100 ng mL^-1^, R&D Systems), and R-spondin-1 (0.5 μg mL^-1^, R&D systems).

Co-culture experiments were performed by transferring Transwell inserts containing 2D-hIO monolayers from the empty plates in which they were grown and differentiated into plates containing differentiated adipocytes. In that case, hIO differentiation medium was maintained in the upper chamber (0.5 mL) but the lower chamber contained the adipocyte differentiation/maintenance medium in which adipocytes had been maintained. At 6-8 hours after beginning co-culture media were changed to serum-free medium (DMEM/F_12_ supplemented with HEPES 15 mM, fatty acid-free BSA 1%, GlutaMAX, and pencillin-streptomycin) for overnight serum starvation where indicated. No co-culture period lasted for more than 24 hours.

### Human induced pluripotent stem cells and mesenchymal progenitor cells

Human induced pluripotent stem cells (hiPSC) were a gift from Dr. Claudia Doege (Columbia University). hiPSC were cultured in mTESR+ (StemCell Technologies) on culture surfaces coated with a thin film of hESC-qualified Matrigel (Corning); culture medium was changed every other day. Cells were split by dissociating to small clusters and replating at a 1:3-1:4 ratio using ReLeSR (StemCell Technologies) once cell clusters reached ∼70% coverage of the culture surface. Medium was supplemented with Y27632 (ROCK inhibitor) 10 μM during and for 24 hours after replating.

hiPSC were differentiated to mesenchymal progenitor cells (MPC) using the STEMdiff Mesenchymal Progenitor Kit (StemCell Technologies), following manufacturer’s instructions. Briefly, hiPSC clusters were dissociated to single cells using GCDR and replated in mTESR+ containing Y27632 (10 μM) at approximately 5 × 10^4^ viable cells cm^-2^ on Matrigel-coated 6-well plates. Two days later, after single cells had proliferated to 70-90% confluence, medium was changed to STEMdiff Mesenchymal Induction Medium; medium was replaced daily for 3 days and then was replaced with MesenCult-ACF Plus medium for 2 days. Cells were then passaged using GCDR and replated on 10-cm dishes coated with Animal Component Free (ACF) Attachment Substrate (StemCell Technologies); all subsequent passages were grown using the same medium and culture substrate. Cells were thereafter passaged once reaching 80% confluence and medium was supplemented with Y-27632 (10 μM) during and for 24 hours replating. MPC differentiation was complete by around 21 days after initial induction. MPC were thereafter passaged upon reaching 80% confluence and used to derive adipocytes.

### Preparation of lentiviral supernatant for MPC infection

Adipocytes were derived from MPC infected with a tetracycline-inducible *PPARG2* lentivirus [17]. All lentiviral constructs were a gift of Dr. Max Friesen (MIT). In order to prepare lentivirus-enriched medium, 293T cells were transfected with plasmids encoding lentiviral structural components (pMDL, pREV, VSVG) and payloads (*PPARG2* or rtTA, at 10 μg per planned MPC plate) using Lipofectamine 3000 (Invitrogen) in low-serum, antibiotic-free Opti-MEM (Gibco). Medium was changed 6 hours later to antibiotic-free DMEM/F_12_ + 10% FBS. At 48 h after infection, viral supernatant (medium) from PPARG2- and rtTA-infected cells were filtered (0.22 μm) and collected. New medium was replaced and then 24 h later was filtered, collected, and added to the previous day’s viral supernatant. In both cases, the presence of lentivirus was confirmed using Lenti-X GoStix (TaKaRa). Viral supernatants were supplemented with a mixture of 60% DMEM/40% FBS and stored at 4ºC if to be used within the next 48 hours or frozen at -80ºC until use. Immediately prior to use in MPC, PPARG2 and rtTA viral supernatants were mixed together at a 1:1 ratio.

### Differentiation of adipocytes

MPC to be differentiated to adipocytes were plated in MesenCult ACF-Plus medium on 12-well dishes coated with ACF Attachment Substrate. Upon reaching confluence, the medium was changed to viral supernatant containing equal proportions of PPARG2 and rtTA supernatant and supplemented with Polybrene (Sigma-Aldrich) at 7.5 μg mL^-1^. Medium was changed 24 h later (Day 0) to adipocyte differentiation medium (ADM) and then changed every other day thereafter. ADM consists of a base medium – DMEM/F_12_ plus Knock-Out Serum Replacement (Gibco) 7.5%, non-essential amino acids (Gibco) 0.5%, and penicillin-streptomycin (Gibco) 1% – supplemented with dexamethasone (1 μM), insulin (10 μg mL^-1^), and rosiglitazone (0.5 μM). Doxycycline was added at 700 ng mL^-1^ to switch on rtTA for the first 16 days of differentiation. Mature adipocytes were used for experiments at or after Day 21.

### Bile acid mixes

Unconjugated bile acid mixes were prepared and dissolved in DMSO according to proportions we have published previously [15]. H10 (*i*.*e*., 10% 12-HBA, 90% non-12-HBA) consists, on a molar basis, of: 7% cholic acid (CA), 3% deoxycholic acid (DCA), 86.85% chenodeoxycholic acid (CDCA), 2.25% ursodeoxycholic acid (UDCA), and 0.90% lithocholic acid (LCA). H90 (*i*.*e*., 90% 12-HBA, 10% non-12-HBA) consists of 63% CA, 27% DCA, 9.65% CDCA, 0.25% UDCA, and 0.10% LCA. BA mixtures were store at -20ºC. All BA treatments were performed by adding BA mixes (or DMSO vehicle) to the culture medium at a final concentration of 100 μM and incubating for 24 hours. FGF-19 secretion into the culture medium following BA treatment was measured by ELISA (Cell Signaling) following manufacturer’s instructions.

### Assays of insulin action

Unless otherwise specified, adipocytes, and hIO, where co-cultured, were thoroughly washed with PBS and then serum-starved overnight in DMEM/F_12_ supplemented with HEPES 15 mM, fatty acid-free BSA 1%, GlutaMAX, and pencillin-streptomycin prior to assays of insulin action.

Lipolysis was measured using the Zen-Bio Human Lipolysis Assay kit, following manufacturer’s instructions. After the overnight serum starvation, for lipolysis experiments, pre-treatment with insulin 100 nM (Humulin-R U-100, Lilly) was added directly to the serum-free medium. After 1 hour of pre-treatment for lipolysis experiments, medium was changed to pre-mixed treatments in lipolysis assay medium: insulin 100 nM (Humulin-R U-100, Lilly) and/or isoproterenol 1 mM (Sigma-Aldrich). Lipolysis was measured over 3 hours, unless otherwise indicated, and assayed colorimetrically according to kit instructions. After the lipolysis measurement period, cells were washed with PBS and lysed in ice-cold RIPA buffer containing PMSF (1 mM, Cell Signaling) and protease/phosphatase inhibitor cocktail (1X, Cell Signaling). Protein quantification was performed by the bicinchoninic acid method (Sigma-Aldrich) for normalization of lipolysis data.

For experiments in which only insulin stimulation was performed (*e*.*g*., for AKT phosphorylation), no pre-treatment step was performed and insulin was applied to cells for 30 min. Cells were then lysed as above and AKT phosphorylation was measured by ELISA (Cell Signaling) following manufacturer’s instructions.

### RT-PCR

RNA was extracted and purified using the RNeasy Mini Kit (QIAgen) for hIO and the RNeasy Lipid Tissue Mini Kit (QIAgen) for adipocytes, in both cases following manufacturer’s instructions. In both cases, on-column DNase treatment was performed using the RNase-Free DNase Set (QIAgen). RNA (1 μg per reaction) was reverse transcribed using the GoScript Reverse Transcription System (Promega). Following reverse transcription, cDNA were diluted 1:10 in sterile water. RT-PCR was then performed using the SYBR Green qPCR Master Mix (Promega). Primer sequences are available upon request. Data were analyzed using the 2^-ΔΔCt^ method after normalization to C_t_ for *TBP*. Unless otherwise noted, all gene-expression data represent normalization of *TBP*-adjusted gene expression to that of the respective experiment’s control condition.

### RNA-seq

RNA was isolated as above. Concentration and purity (RIN ≥ 8.0) were verified by the Columbia University Molecular Pathology core facility using the Agilent Bioanalyzer 2100. RNA (2 μg per sample) were then submitted for RNA-seq to be performed by the J.P. Sulzberger Columbia Genome Center using TruSeq Stranded mRNA Library Prep Kit (Illumina) for library preparation followed by sequencing using NovaSeq (2×100 bp). Data analysis was performed in conjunction with the Genome Center.

### Statistical analysis

All quantitative data are expressed as mean ± SEM. Unless otherwise noted, statistical analyses were performed as ANOVA followed by Tukey’s *post hoc* test for multiple comparisons; one- and two-way ANOVA were used for comparisons involving one or two independent variable(s), respectively. Unpaired, two-tailed student’s *t* test was used for experiments comparing only two conditions. Significance cutoff was set at *p* < 0.05 for all experiments.

## Results

### Human intestinal organoids

We derived human intestinal organoids (hIO) from duodenal biopsy specimens taken during endoscopies performed for other purposes. Culture of hIO in Matrigel domes resulted in the development of three-dimensional (3D-hIO) cystic structures over the course of 21 days (Figure 1A). Given the wide array of protocols available for culture and differentiation of hIO [18], we sought to make comparisons between two of the more widely used methods. We tested homemade (see Methods) (CCM, left column) [16] and preformulated (SCT-ODM, right column) hIO cocktails and found that both culture methods produced similar morphologic changes during the differentiation process (Figure 1B). Gene expression profiling of hIO at day 21 of differentiation demonstrated the presence of genes typical of enterocytes (*ANPEP, FGF19, IBABP, VIL1*), goblet cells (*ITF*), and Paneth cells (*LYZ1*) (Figure 1C), but enteroendocrine cell markers (*CHGA, GIP, GCG*) were not detectable in either case (data not shown).

**Figure 1.**
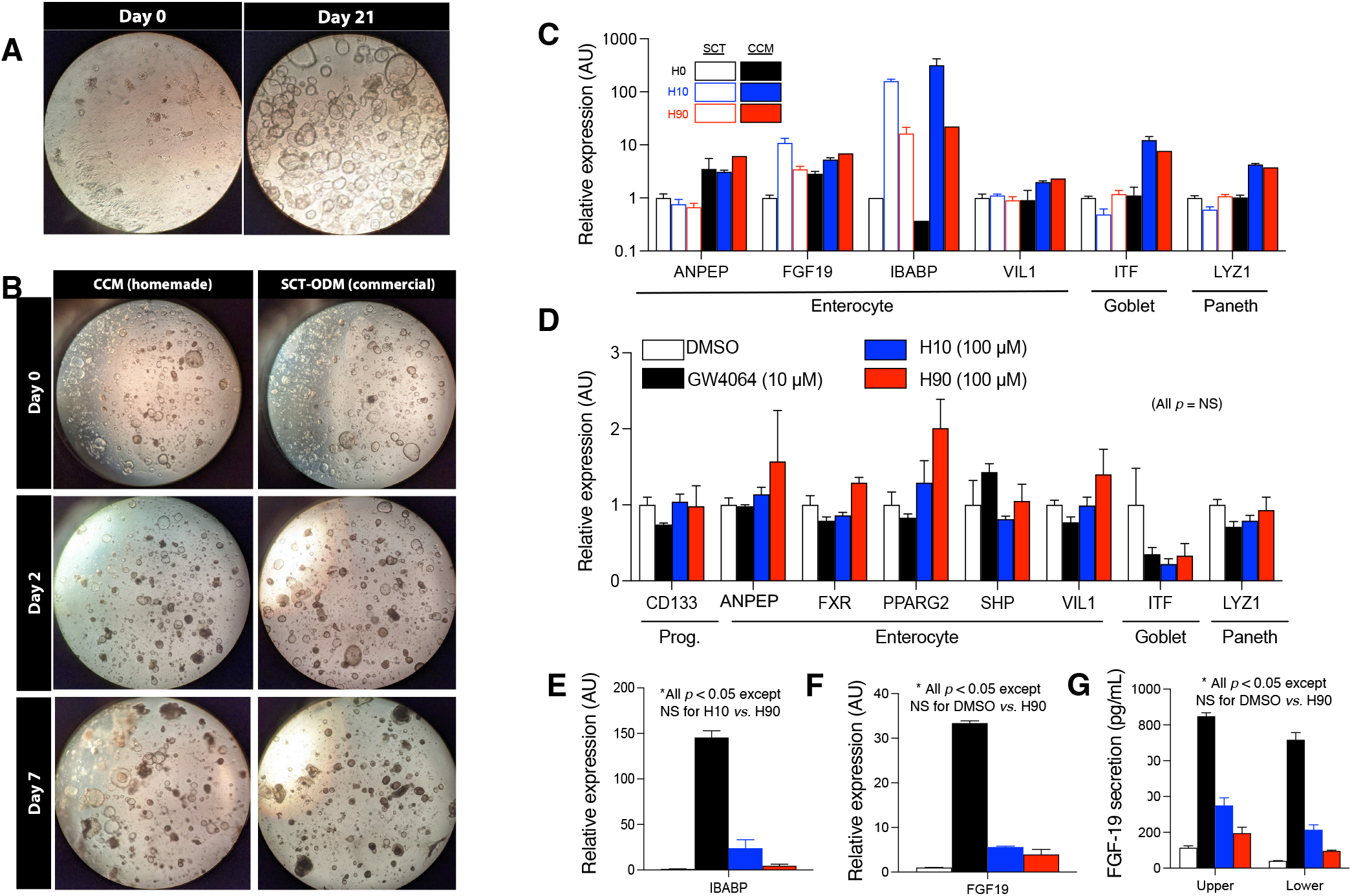
Derivation and characterization of primary human intestinal organoids (hIO). (A) hIO were derived from duodenal biopsy samples and plated in Matrigel domes on Day 0 (left panel) and by Day 21 (right panel) had proliferated as three-dimensional immature spheroids. (B) Comparison of two hIO culture methods across 7 days of differentiation from spheroids to mature hIO: homemade “CCM” medium (Crypt Culture Medium, left panel) *versus* “SCT-ODM” (StemCell Technologies Organoid Differentiation Medium, right panel). (C) Comparison of the expression of key 3D-hIO genes (by RT-PCR) in response to 100-μM mixes of varied 12-HBA composition (H0, black; H10, blue; H90, red), stratified by differentiation in SCT-ODM (empty bars) *versus* CCM (filled bars). (D-F) Characterization of gene expression in SCT-ODM-cultured hIO grown in 2D monolayer in response to FXR agonist (GW4064, black) or bile acid mixes (H10, blue; H90, red). In all experiments, “Relative expression” indicates normalization of *TBP*-adjusted gene expression to that of the control condition. AU = arbitrary units. (G) ELISA-based quantification of FGF-19 levels secreted into the medium of either the upper (left group) or lower (right group) chamber of the Transwell setup.

We next evaluated the functional suitability of each method for studying the effects of BA mixes on gene expression. For this, we produced unconjugated BA mixes of different 12-HBA content; H10 denotes 10% 12-HBA (*i*.*e*., 90% non-12-HBA), and H90 denotes 90% 12-HBA. Based on our group’s previous work [15], we expect robust induction of *FGF19* expression by H10 compared to H0 (vehicle) and, to a lesser extent, by H90. We found that these expected changes in *FGF19* expression were reproduced with the commercial formulation but not with the homemade medium. As *IBABP* is preferentially expressed in the ileum [19], its baseline expression in these duodenum-derived hIO was near the lower limit of detection, leaving us unable to make reliable comparisons in the H0 condition. However, *IBABP* expression, like *FGF19*, was robustly induced by H10 and, to a lesser extent, by H90. As the commercial formulation appeared to yield results more consistent with published findings [15], we favored it to perform the experiments in 2D-hIO below.

In preparation for adipocyte co-culture experiments, 3D-hIO were dissociated and replated as a monolayer (2D-hIO) on semipermeable membranes in Corning Transwell dishes. 2D-hIO grew to confluence over the course of about a week but, based on our observation of FGF-19 accumulation in both the upper (apical) and lower (basolateral) compartments, the monolayer did not achieve impermeability. Nevertheless, 2D-hIO robustly expressed genes typical of enterocytes (*e*.*g*., *ANPEP, VIL1*), goblet cells (*ITF*), and Paneth cells (*LYZ1*), in addition to retaining expression of *CD133*, typical of intestinal progenitor cells (Figure 1D) [20]. Interestingly, 2D-hIO baseline expression of *IBABP* was, unlike in 3D-hIO, well within the range of detection (Figure 1E). However, we again were unable to detect expression of enteroendocrine cell markers such as *CHGA* or *GIP*.

Having established 2D-hIO cultures, we turned to studying the impact of BA signaling on their gene expression profiles. Treatment for 24 h with BA mixes (100 μM of H10 or H90) or, as a positive control, the FXR agonist GW4064 resulted in strong induction of the expression of *IBABP* (Figure 1E) and *FGF19* (Figure 1F), reflected as well in FGF-19 protein secreted into the medium (Figure 1G). In each case, H10 induced a larger effect than H90, consistent with our prior observations [15]. BA mixes and GW4064 did not affect the expression of other enterocyte genes tested, nor of goblet or Paneth cell markers (Figure 1B). In order to appreciate broader changes in gene expression patterns with BA treatment, we performed RNA-seq analysis on 2D-hIO cultured for 24 hours with H0 (DMSO), H10, or H90. As illustrated in Table 1, treatment with H10 significantly (P_adj_<0.05) altered 63 genes compared to H0 and 144 genes compared to H90, while H90 significantly altered 974 genes compared to H0. True to form, the BA transporter *SLC51A* (OSTα) placed among the top ten most significantly altered genes with H10 and H90, while *FGF19* was near the top of the list for H10 compared both to H0 and to H90. Given the negative impact specifically of 12-HBA on insulin sensitivity [6, 13], we are particularly interested in the H90 *vs*. H0 comparison. DAVID pathway analysis was notable for highly significant changes in several metabolically relevant processes, including cholesterol biosynthesis, cellular response to hypoxia, and cellular response to insulin. The list of the most significantly altered genes features entries relevant to each of these pathways, including *SQLE* (squalene monooxygenase), *RHOB* (RAS homolog member B), and *INSIG1* (insulin-induced gene 1), respectively, within the top ten.

**Table 1.** RNA-seq of 2D-hIO cultured with bile acid mixes. 2D-hIO were treated for 24 hours with bile acid mixes (100 μM) of varying 12-HBA content: H0, H10, or H90. Depicted in the table are comparisons of each condition, including: (i) the number of genes whose adjusted *p* value was < 0.05, (ii) the top ten most highly altered biological processes according to DAVID gene-ontology analysis, and (iii) the top ten most highly altered individual genes.

| Comparison | Overall<br># genes | DAVID analysis |  |  | Top genes |  |
| --- | --- | --- | --- | --- | --- | --- |
|  |  | Pathway | # genes | p value | Gene | Name |
| H10 vs. H0 | 63 | Cell cycle | 11 | 2.25 × 10<sup>-5</sup> | SLC51A | Organic solute transporter α |
|  |  | Acetylation | 22 | 7.12 × 10<sup>-4</sup> | RRM2 | RNR regulatory subunit M2 |
|  |  | ATP binding | 13 | 7.95 × 10<sup>-4</sup> | ANLN | Anillin |
|  |  | Sterol metabolism | 4 | 0.001 | CHAF1A | Chromatin assembly factor 1A |
|  |  | Cell division | 7 | 0.001 | FGF19 | Fibroblast growth factor 19 |
|  |  | Phosphoprotein | 38 | 0.001 | CALD1 | Caldesmon 1 |
|  |  | Polymorphism | 49 | 0.001 | MKI67 | Ki-67 |
|  |  | Ubl conjugation | 14 | 0.001 | TGM2 | Tissue transglutaminase |
|  |  | Lipid metabolism | 7 | 0.002 | AOX1 | Aldehyde oxidase 1 |
|  |  | Isopeptide bond | 11 | 0.002 | OAS3 | 2'5'-oligoadenylate synthase 3 |
| H90 vs. H0 | 974 | Cholesterol biosynthesis | 19 | 6.28 × 10<sup>-14</sup> | RHOB | RAS homolog member B |
|  |  | Cellular response to hypoxia | 18 | 6.72 × 10<sup>-6</sup> | CCL20 | Chemokine ligand 20 |
|  |  | Ephrin receptor signaling | 16 | 2.79 × 10<sup>-5</sup> | SPNS2 | Sphingolipid transporter 2 |
|  |  | Isoprenoid biosynthesis | 7 | 3.74 × 10<sup>-5</sup> | BIRC3 | Baculoviral IAP repeat containing 3 |
|  |  | Apoptotic process | 52 | 6.74 × 10<sup>-5</sup> | KLF2 | Krüppel-like factor 2 |
|  |  | Cellular response to insulin | 14 | 1.33 × 10<sup>-4</sup> | CD24 | CD24 sialoglycoprotein |
|  |  | Negative caspase regulation | 13 | 1.79 × 10<sup>-4</sup> | INSIG1 | Insulin-induced gene 1 |
|  |  | Rho signal transduction | 11 | 1.87 × 10<sup>-4</sup> | SLC51A | Organic solute transporter α |
|  |  | Protein stabilization | 19 | 2.00 × 10<sup>-4</sup> | SDC1 | Syndecan 1 |
|  |  | Negative apoptotic regulation | 42 | 3.28 × 10<sup>-4</sup> | SQLE | Squalene monooxygenase |
| H90 vs. H10 | 144 | Mitotic cytokinesis | 6 | 2.00 × 10<sup>-6</sup> | ANLN | Anillin |
|  |  | Nuclear division | 11 | 1.43 × 10<sup>-5</sup> | SLC51A | Organic solute transporter α |
|  |  | Sister chromatid cohesion | 7 | 1.09 × 10<sup>-4</sup> | RRM2 | RNR regulatory subunit M2 |
|  |  | Chromosome segregation | 6 | 1.40 × 10<sup>-4</sup> | H4C15 | H4 clustered histone 15 |
|  |  | Cell division | 11 | 2.54 × 10<sup>-4</sup> | FGF19 | Fibroblast growth factor 19 |
|  |  | Cell proliferation | 11 | 3.62 × 10<sup>-4</sup> | EXO1 | Exonuclease 1 |
|  |  | G<sub>2</sub>/M transition | 7 | 5.14 × 10<sup>-4</sup> | KIF11 | Kinesin family member 11 |
|  |  | Mitotic spindle assembly | 4 | 0.002 | ITGB1P1 | Integrin β<sub>1</sub> pseudogene 1 |
|  |  | DNA damage response | 7 | 0.004 | TOP2A | Topoisomerase 2α |
|  |  | DNA replication | 6 | 0.006 | ABCB4 | Multidrug resistance protein 3 |

### iPS-derived adipocytes

Adipocytes were derived from human iPS cells though inducible lentiviral overexpression of *PPARG2* after initial differentiation to mesenchymal progenitor cells (MPC) [17]. After a 14-day incubation in adipogenesis medium, MPCs differentiated to adipocytes at high efficiency, both morphologically (Figure 2A) and in the expression of classic adipocyte genes such as *ADIPOQ, AP2, FASN*, and *LPL* (Figure 2B); we verified adiponectin production and secretion into the culture medium by ELISA (at least 1 μg per mg cellular protein). Interestingly, the adipocytes also express uncoupling protein-1 (*UCP1*), typical of brown or beige adipocytes. We next confirmed that iPS-derived adipocytes are capable of lipolysis, measured as glycerol release, that is inducible with the pan-β-adrenergic agonist isoproterenol (1 μM) and inhibited by insulin (100 nM) (Figure 1C). We further demonstrate the insulin responsiveness of the cells by demonstrating induction of AKT phosphorylation, 30 minutes after acute insulin treatment (Figure 2D). Because of the presence of insulin and other growth factors in the maintenance culture medium, acute induction of AKT phosphorylation with insulin was much more clearly elicited with overnight serum starvation prior to insulin treatment.

**Figure 2.**
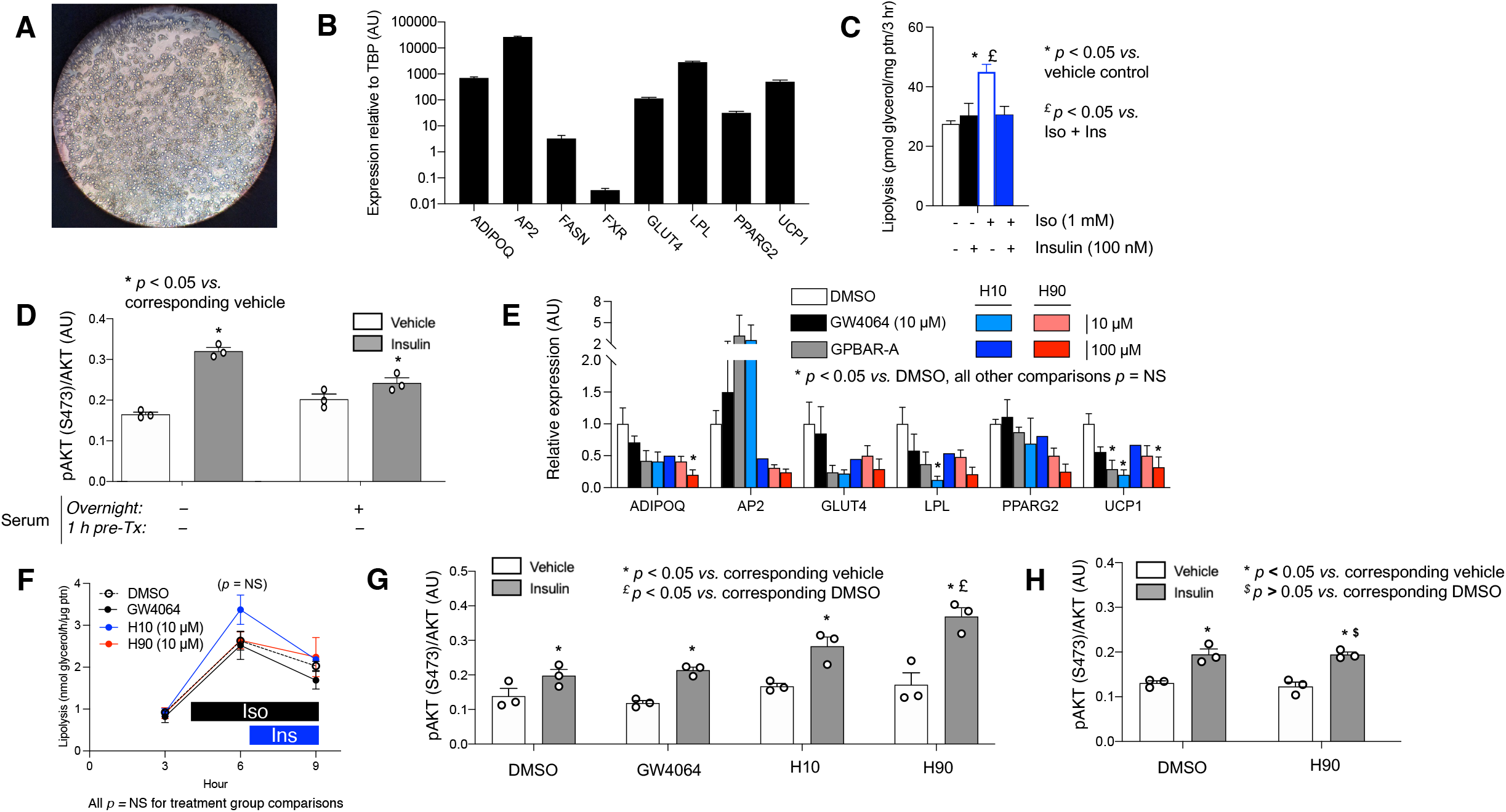
Characterization of hiPSC-derived adipocytes. (A) hiPSC-derived adipocytes in culture following the 21-day differentiation protocol. (B) Expression of key adipocyte genes by RT-PCR, normalized to expression of *TBP*. (C) Assessment of adipocyte lipolysis (*i*.*e*., glycerol release) over 3 hours in response to isoproterenol (1 mM) and/or insulin (100 nM). (D) ELISA assessment of AKT (S473) phosphorylation in response to vehicle (white) or insulin (100 nM, gray), with (left group) or without (right group) preceding overnight serum. (E) RT-PCR-based gene expression in adipocytes in response to 24-hour treatment with DMSO (white), FXR agonist (GW4064, black), TGR5 agonist (GPBAR-A, gray), or bile acid mixes (H10, blue; H90, red). Bile acid mixes were used at both lower (10 μM, lighter coloration) and higher concentrations (100 μM, darker coloration). “Relative expression” indicates normalization of *TBP*-adjusted gene expression to that of the control condition. AU = arbitrary units. (F) Change in rate of adipocyte lipolysis in response to serial treatments with isoproterenol (Iso, 1 nM) ± insulin (Ins, 100 nM) as a function of treatment with DMSO (white), GW4064 (black), H10 (blue, 10 μM), or H90 (red, 10 μM). (G-H) ELISA assessment of AKT (S473) phosphorylation in response to vehicle (white) or insulin (100 nM, gray) following 24-hour treatment with GW4064 (10 μM), H10 (10 μM), or H90 (10 μM). Statistical analysis done as noted in the figure; * *p* < 0.05 by one-way ANOVA and Tukey’s *post hoc* test for multiple comparisons.

Prior to introducing adipocytes to hIO, we sought to understand the impact of BA treatment on adipocytes *per se*. Treatment with H10, H90, GW4064, or TGR5/GPBAR-A agonist generally decreased expression of classic adipocyte genes, but these changes only reached statistical significance in scattered cases (Figure 2E). We do not have quantitative data on the effect of BA on adipocyte viability, but there was no qualitative differences between treatments on microscopic inspection. Moreover, neither treatment with the BA mixes or with GW4064 significantly altered the stimulation of lipolysis by isoproterenol or insulin’s inhibition thereof (Figure 2F). Similarly, there was no systematic effect of BA or FXR agonist treatment on phosphorylation of AKT in response to acute insulin treatment (Figure 2G). Although there appeared to be a trend toward increased AKT pS473 following treatment with H90, this effect was not reproducible (Figure 2H). Thus, on the whole, BA did not appear to directly regulate adipocyte identity or insulin signaling in the limited assays we conducted.

We further explored the impact of BA on adipocyte gene expression by RNA-seq (Table 2). Few genes’ expression were altered by treatment with H10, whether in comparison to H0 (18 genes) or to H90 (5 genes). Yet, treatment with H90 altered expression of 180 genes *versus* H0, suggesting a specific role for 12-HBA in regulating adipocyte gene expression. DAVID gene-ontology analysis indicated that these genes are most prominently involved in DNA replication and repair. However, certain metabolically relevant genes were also significantly upregulated, including *MLXIPL* (carbohydrate-responsive element binding protein, ChREBP) and *CAV1* (caveolin-1), the latter of which has been implicated in the control of systemic glucose metabolism by adipocytes [21].

**Table 2.** RNA-seq of adipocytes cultured with bile acid mixes. hiPSC-derived adipocytes were treated for 24 hours with bile acid mixes (100 μM) of varying 12-HBA content: H0, H10, or H90. Depicted in the table are comparisons of each condition, including: (i) the number of genes whose adjusted *p* value was < 0.05, the top ten most highly altered biological processes according to DAVID gene-ontology analysis, and the top ten most highly altered individual genes.

| Comparison | Overall # genes | DAVID analysis |  |  | Top genes |  |
| --- | --- | --- | --- | --- | --- | --- |
|  |  | Pathway | # genes | p value | Gene | Name |
| H10 vs. H0 | 18 | Data inconclusive due to small number of genes |  |  | SERPINE1 | Plasminogen activator-inhibitor 1 |
|  |  |  |  |  | XRN1 | 5'-3' exoribonuclease 1 |
| | | | | | POLG | DNA polymerase $\gamma$ |
|  |  |  |  |  | CDC6 | Cell division cycle 6 |
|  |  |  |  |  | ANPEP | Aminopeptidase N |
|  |  |  |  |  | GREM1 | Gremlin 1 |
|  |  |  |  |  | ATF7 | cAMP-activating transc. factor 7 |
|  |  |  |  |  | CADM1 | Cell adhesion molecule 1 |
|  |  |  |  |  | HMGA1 | High-mobility group protein I/Y |
|  |  |  |  |  | UBE2F-SCLY | UBE2F-SCLY readthrough |
| H90 vs. H0 | 180 | DNA unwinding in replication | 8 | $7.3 \times 10^{-10}$ | SERPINE1 | Plasminogen activator-inhibitor 1 |
| | | DNA replication initiation | 8 | $3.4 \times 10^{-9}$ | HMGA1 | High-mobility group protein I/Y |
| | | Double-strand break repair | 6 | $3.5 \times 10^{-8}$ | ANPEP | Aminopeptidase N |
| | | DNA replication | 11 | $1.0 \times 10^{-7}$ | LAMC2 | Laminin subunit $\gamma 2$ |
| | | Pre-replicative complex assembly | 5 | $3.8 \times 10^{-7}$ | CDC6 | Cell division cycle 6 |
| | | Cell cycle | 16 | $6.0 \times 10^{-7}$ | S100A10 | S100 Ca-binding protein A10 |
| | | Mitotic cell cycle | 11 | $9.9 \times 10^{-7}$ | MCM2 | DNA replication licensing factor MCM2 |
| | | DNA repair | 11 | $2.6 \times 10^{-4}$ | UHRF1 | Ubiquitin-like w/ PHD and RNF dom. 1 |
| | | Aging | 9 | $3.0 \times 10^{-4}$ | IGFBP6 | IGF binding protein 6 |
| | | Angiogenesis | 10 | $3.7 \times 10^{-4}$ | RRM2 | Ribonucleotide reductase M2 |
| H90 vs. H10 | 5 | Data inconclusive due to small number of genes |  |  | PRG4 | Proteoglycan 4 |
|  |  |  |  |  | HARS2 | His-tRNA synthetase 2, mitochond. |
|  |  |  |  |  | RHEBP1 | RHEB pseudogene 1 |
|  |  |  |  |  | PMS2P8 | PMS1 homologue 2 pseudogene 8 |
| | | | | | Remainder with $p_{adj} > 0.05$ | |

### Co-culture experiments

After having established some baseline characteristics of its individual components, we moved to implement the hIO/adipocyte co-culture system. As illustrated in Figure 3A, 3D-hIO were plated in monolayer atop semipermeable membranes in Transwells that were then suspended above a monolayer of adipocytes plated in the wells of cell culture dishes for 24 hours in each experiment. Once the system was established, we first sought the impact of co-culture on lipid metabolism. Co-culture had no apparent effect on total TG content of adipocytes (Figure 3B). Interestingly, hIO were also found also to contain appreciable quantities of stored TG [22], representing about 10% of the adipocyte TG content when normalized to total cellular protein. Neither isoproterenol nor albumin-bound palmitate treatment altered intracellular triglyceride levels in either compartment.

**Figure 3.**
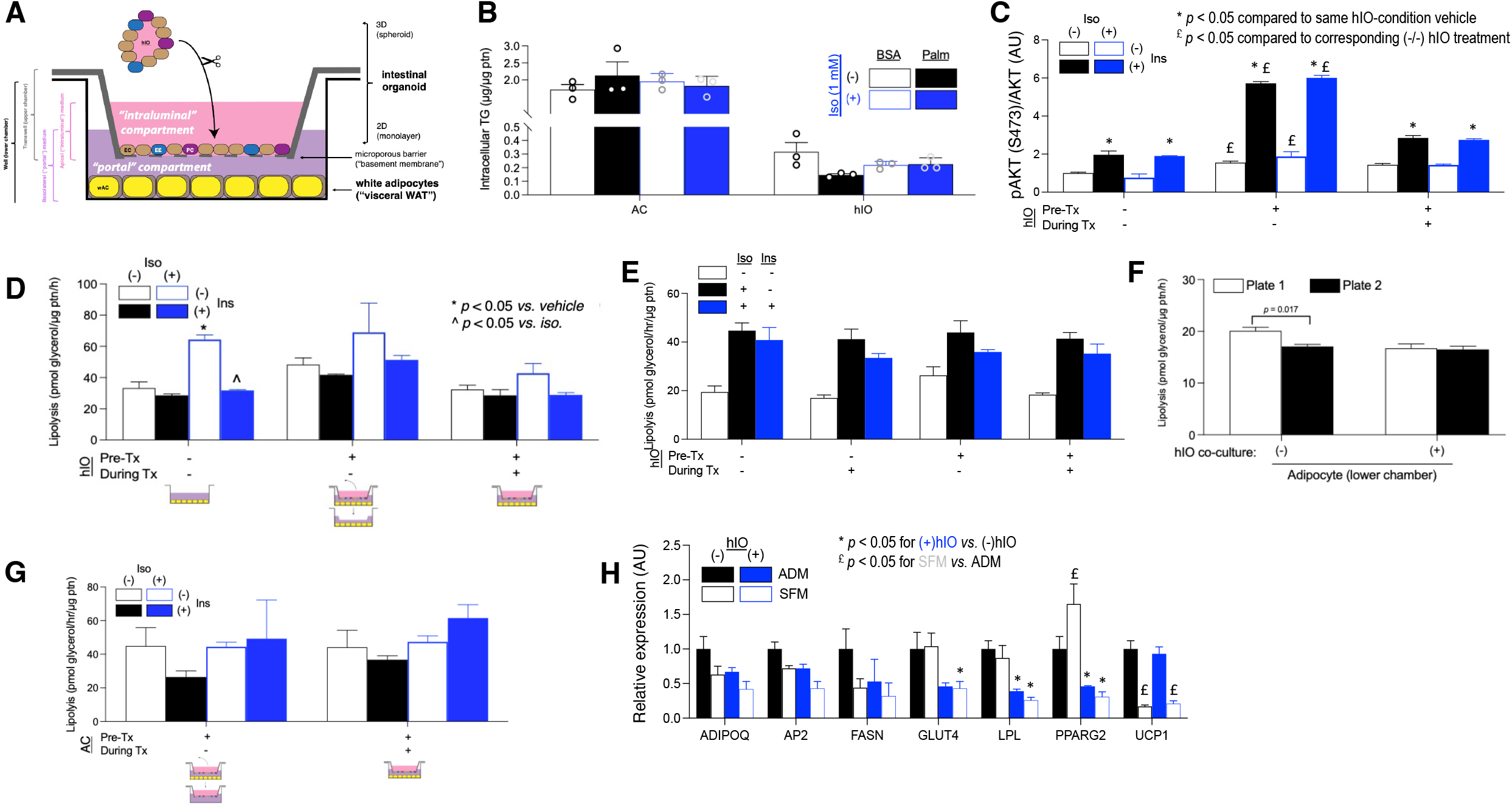
Co-culture of 2D-hIO and hiPSC-derived adipocytes. (A) Schematic of co-culture setup in Corning Transwell plates. (B) Intracellular triglyceride (TG) content in adipocytes (AC, left group) or 2D-hIO (right group) in response to overnight culture with BSA (open bars) or BSA-conjugated palmitate (Palm, filled bars) and acute (3-hour) treatment without (black) or with (blue) isoproterenol (Iso, 1 mM). (C) ELISA-based AKT (S473) phosphorylation and (D-E) lipolysis (glycerol release) in adipocytes in response to insulin (Ins, 100 nM, black) and/or isoproterenol (Iso, 1 mM, blue), as a function of the absence of hIO (left group) or presence of hIO for 24 hours before (right two groups) ± during (right group) acute treatments. (F) Basal adipocyte lipolysis (glycerol release) of “sister” plates in the absence (left group) or presence (right group) of hIO. (G) Lipolysis by 2D-hIO in same experiment as panel *D*. (H) Gene expression by RT-PCR in adipocytes cultured in adipocyte differentiation medium (ADM, filled bars) or serum-free medium (SFM, empty bars), either in the absence (black) or presence (blue) of 2D-hIO. Statistical analysis done as noted in the figure by (C, D, G, H) two-way ANOVA and Tukey’s *post hoc* test for multiple comparisons or (F) unpaired, two-tailed student’s *t* test.

We next explored the effect of co-culture on insulin-responsive processes. We found that 24-hour co-culture with hIO significantly enhanced AKT phosphorylation in adipocytes in response to insulin if hIO were removed prior to insulin stimulation but this effect was attenuated if hIO remained present during the insulin stimulation (Figure 3C). Co-culture appeared to blunt the induction of lipolysis by isoproterenol, regardless of whether hIO remained present during the 3-hour lipolysis assessment period, but did not significantly affect basal or insulin-treated lipolysis (Figure 3D). However, this finding was not consistently reproducible, failing to replicate in one of three independent experiments (Figure 3E). There are three possible explanations for this observation. First, hIO were derived from a different human donor for the third experiment *versus* the first two. We question the importance of a putative co-culture effect on lipolysis that is not sufficiently robust to replicate across individuals. Second, even two “sister” plates – that is, differentiated at the same time, from a single parent plate, and treated simultaneously – differ in rate of basal lipolysis (Figure 3F). Thus, differing plate layouts likely led to divergent results for the same experiment, again demonstrating the limitations of these non-immortalized cell models for lipolysis assays. Finally, lipolysis by the hIO themselves may confound interpretation of adipocyte lipolysis. We demonstrated that glycerol accumulated in the upper-chamber culture medium even after separating the 2D-hIO monolayer from the adipocytes and washing it extensively with PBS (Figure 3G). It also did not respond to isoproterenol or insulin and so differentially regulated hIO lipolysis may have confounded the interpretation of adipocyte lipolysis in co-culture. Thus, overall, we conclude that these stem cell-based systems are highly variable in co-culture lipolysis assays, and large numbers of replicates would be required to draw strong conclusions.

Finally, we investigated changes in adipocyte gene expression occurring in response to 24 hours of co-culture (Figure 3H). In order to mimic the effects of the fed *versus* fasted states, we further stratified the analysis by treatment in adipocyte differentiation medium (ADM) or serum-free medium (SFM), respectively. Overall, we identified three gene-expression patterns among the genes tested. In the first group, both serum starvation and hIO tended to decreased gene expression, perhaps synergistically: *ADIPOQ, AP2, FASN*. In the second, hIO decreased expression of genes otherwise not decreased by serum starvation: *SLC2A4* [GLUT4], *LPL, PPARG2*). Finally, the only gene tested that was clearly unaffected by hIO co-culture was *UCP1*, which was also the most strongly affected by serum starvation.

### Transcriptomic data

We performed RNA-seq analysis on adipocytes and 2D-hIO, each both alone and in co-culture with the other for 24 hours, in order to take an unbiased approach to the functional significance of their interactions. As illustrated in Figure 4A, all of the 25 most significantly altered adipocyte genes were upregulated by the presence of hIO. Gene-ontology analysis indicates that the most highly affected adipocyte processes include the hypoxia response and anaerobic glucose metabolism (Figure 4C). On the other hand, hIO gene expression was more variably affected by the presence of adipocytes, featuring similar enrichment with downregulated and upregulated genes among the top 25 (Figure 4B). Overall, the most highly significant changes occur in upregulated genes, largely represented by lipid-(and particularly sterol-) metabolic processes (Figure 4C).

**Figure 4.**
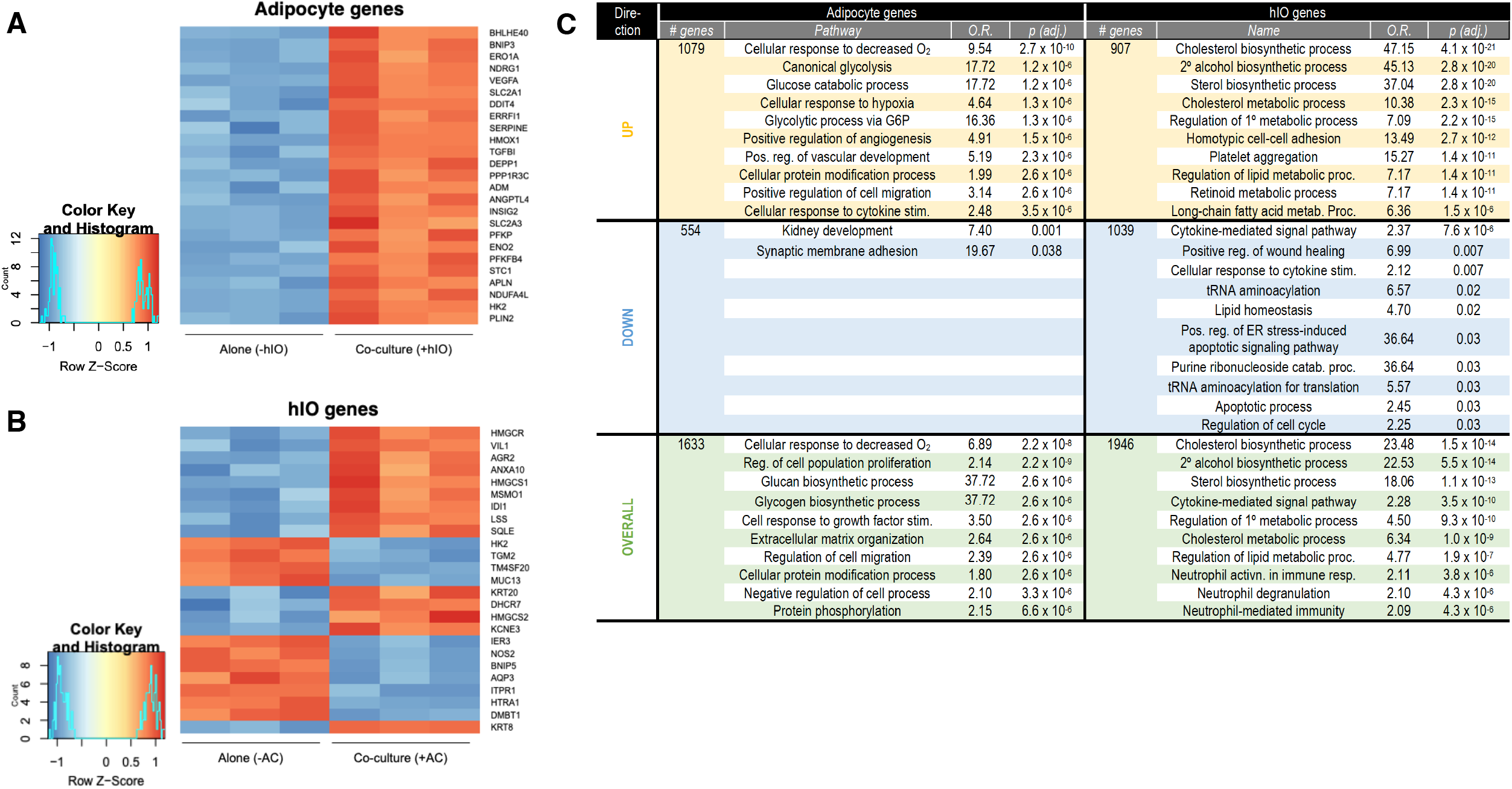
RNA-seq of co-cultured hiPSC-derived adipocytes and 2D-hIO. RNA-seq was performed in (A) adipocytes (AC) and (B) 2D-hIO that were cultured either alone or in co-culture. Heat maps indicate the most significantly altered genes in each condition; red indicates increased expression *versus* the alternate culture condition while decreased expression is in blue. (C) Enrichr-based pathway analysis for each cell-type comparison based on those that are upregulated (yellow) or downregulated (blue), or based on absolute value of change (green). Only processes with *p*_adj_ < 0.05 are shown.

## Discussion

In this report, we have presented pilot data from a co-culture system that uses human induced pluripotent stem cell-derived adipocytes and primary human duodenal organoids to model interactions, particularly as driven by bile acids, between the intestinal epithelium and visceral white adipose tissue. First, we demonstrate that primary 2D-hIO respond to treatment with BA mixtures of varied 12-HBA content in a manner consistent with our prior studies. Moreover, we have generated transcriptomic data indicating that 12-HBA differentially regulate the expression of genes involved in intestinal sterol metabolism and the response to hypoxia. Next, we demonstrated that there is no clear effect of BA treatment on the expression of classic adipocyte genes or on key insulin-regulated processes in adipocytes. However, there may be a specific impact of 12-HBA on expression of key metabolism-associated genes in adipocytes, such as those encoding ChREBP [23] or CAV1 [21]. Putting the two systems together, we found co-culture of adipocytes and hIO did not affect adipocyte lipolysis in a consistent way but did decrease expression of key adipocyte genes. In fact, co-culture exerted large-scale transcriptomic changes on gene expression in both cellular compartments: the presence of hIO most significantly ramped up adipocyte expression of genes involved in the response to hypoxia and anaerobic glucose metabolism, while adipocytes, in turn, most strongly stimulated expression of hIO genes involved in lipid metabolism.

Because the primary purpose of this report is to provide interim guidance on the optimization of a metabolically relevant co-culture system, the experiments we present have clear limitations. We have found substantial plate-to-plate variability in cellular phenotypes, even when derived and cultured in parallel. For example, even “sister” plates of adipocytes can exhibit significantly different basal lipolytic rates. As such, the results of lipolysis studies can vary depending on the experimental design *vis-à-vis* plate layout. We therefore urge caution in the design and interpretation of such experiments.

Nevertheless, we believe that our preliminary findings have important implications for our understanding of how the intestine informs the “visceralization” of associated adipose tissue. Most notably, hIO influence adipocyte gene-expression profiles and vice-versa. Adipocytes increase expression of genes related to the hypoxia response in the presence of hIO. This may help to explain the observation that the portal blood supply of mesenteric WAT is largely responsible for its dysmetabolic qualities [3]. Concordantly, the intestinal epithelium has been shown to influence the hypoxia response in mesenteric WAT, affecting both adipocyte lipid metabolism and gene expression [24]. This is potentially significant because the inflammatory response to the hypoxic tissue microenvironment has been invoked to explain the particular negative impact of visceral WAT on insulin sensitivity [25, 26]. Suggestively, expression of several classic adipocyte genes is lower in hIO-exposed adipocytes, as in visceral WAT from severely obese, insulin-resistant subjects [27]. Yet, by more direct measures, we did not detect any apparent decrement in insulin action in adipocytes exposed to hIO; in fact, AKT phosphorylation may even be enhanced. Transient or mild hypoxia may have an insulin-sensitizing effect on adipocytes [28, 29], and this should be explored in further experiments in our system. However, we also cannot rule out the possibility that the presence of additional cells of any kind in the upper chamber might incite a relative hypoxic state due to competition for oxygen in the culture medium.

The other major transcriptomic change we have noted to arise from co-culture is strong induction of gene expression related to lipid – and especially sterol – metabolism in hIO. Indeed, among the most highly upregulated genes in hIO following adipocyte co-culture are such canonical cholesterol biosynthetic genes as *HMGCR* (HMG-CoA reductase), *SQLE* (squalene monooxygenase), and *LSS* (lanosterol synthase). Enterocytes occupy a unique position in the regulation of sterol metabolism, balancing uptake of dietary sterols with *de novo* sterol synthesis and disposal *via* transintestinal cholesterol excretion (TICE). Constitutive activation of sterol synthesis in the intestine by enterocyte-specific knockout of the key sterol-sensing negative-feedback mediators *Insig1* and *Insig2* results in cholesterol accumulation in intestine, liver, and blood [30]. Conversely, enterocyte-specific knockout of *Srebp2* impairs *de novo* cholesterol synthesis, resulting in complete dependence on dietary cholesterol to prevent lethal enteropathy [31]. Intestine-specific knockout of the leptin receptor has been shown to significantly increase intestinal cholesterol content [32], suggesting a role for adipocytes in the regulation of intestinal cholesterol metabolism. However, to our knowledge, no studies have directly addressed this question, particularly as regards *de novo* sterol synthesis *per se*. Given the implications of intestinal sterol handling for cardiovascular disease, this may prove to be a fruitful avenue for ongoing research using this co-culture system.

